# The Multiple Sclerosis Genomic Map: Role of peripheral immune cells and resident microglia in susceptibility

**DOI:** 10.1101/143933

**Authors:** International Multiple Sclerosis Genetics Consortium, NA Patsopoulos, SE Baranzini, A Santaniello, P Shoostari, C Cotsapas, G Wong, AH Beecham, T James, J Replogle, I Vlachos, C McCabe, T Pers, A Brandes, C White, B Keenan, M Cimpean, P Winn, IP Panteliadis, A Robbins, TFM Andlauer, O Zarzycki, B Dubois, A Goris, H Bach Sondergaard, F Sellebjerg, P Soelberg Sorensen, H Ullum, L Wegner Thoerner, J Saarela, I Cournu Rebeix, V Damotte, B Fontaine, L Guillot Noel, M Lathrop, S Vukusik, A Berthele, V Biberacher, D Buck, C Gasperi, C Graetz, V Grummel, B Hemmer, M Hoshi, B Knier, T Korn, CM Lill, F Luessi, M Mühlau, F Zipp, E Dardiotis, C Agliardi, A Amoroso, N Barizzone, MD Benedetti, L Bernardinelli, P Cavalla, F Clarelli, G Comi, D Cusi, F Esposito, L Ferrè, D Galimberti, C Guaschino, MA Leone, V Martinelli, L Moiola, M Salvetti, M Sorosina, D Vecchio, A Zauli, S Santoro, M Zuccalà, J Mescheriakova, C van Duijn, SD Bos, EG Celius, A Spurkland, M Comabella, X Montalban, L Alfredsson, I Bomfim, D Gomez-Cabrero, J Hillert, M Jagodic, M Lindén, F Piehl, I Jelčić, R Martin, M Sospedra, A Baker, M Ban, C Hawkins, P Hysi, S Kalra, F Karpe, J Khadake, G Lachance, P Molyneux, M Neville, J Thorpe, E Bradshaw, SJ Caillier, P Calabresi, BAC Cree, A Cross, M Davis, PWI de Bakker, S Delgado, M Dembele, K Edwards, K Fitzgerald, IY Frohlich, PA Gourraud, JL Haines, H Hakonarson, D Kimbrough, N Isobe, I Konidari, E Lathi, MH Lee, T Li, D An, A Zimmer, A Lo, L Madireddy, CP Manrique, M Mitrovic, M Olah, E Patrick, MA Pericak-Vance, L Piccio, C Schaefer, H Weiner, K Lage, A Compston, D Hafler, HF Harbo, SL Hauser, G Stewart, S D’Alfonso, G Hadjigeorgiou, B Taylor, LF Barcellos, D Booth, R Hintzen, I Kockum, F Martinelli-Boneschi, JL McCauley, JR Oksenberg, A Oturai, S Sawcer, AJ Ivinson, T Olsson, PL De Jager, Murray Barclay, Laurent Peyrin-Biroulet, Mathias Chamaillard, Jean-Frederick Colombe, Mario Cottone, Anthony Croft, Renata D’Incà, Jonas Halfvarson, Katherine Hanigan, Paul Henderson, Jean-Pierre Hugot, Amir Karban, Nicholas A Kennedy, Mohammed Azam Khan, Marc Lémann, Arie Levine, Dunecan Massey, Monica Milla, Grant W Montgomery, Sok Meng Evelyn Ng, Ioannis Oikonomou, Harald Peeters, Deborah D. Proctor, Jean-Francois Rahier, Rebecca Roberts, Paul Rutgeerts, Frank Seibold, Laura Stronati, Kirstin M Taylor, Leif Törkvist, Kullak Ublick, Johan Van Limbergen, Andre Van Gossum, Morten H. Vatn, Hu Zhang, Wei Zhang, Australia and New Zealand IBDGC, Belgium Genetic Consortium, Initiative on Crohn and Colitis, NIDDK IBDGC, United Kingdom IBDGC, Wellcome Trust Case Control Consortium

**Author notes:** Correspondence to: Philip L. De Jager, MD PhD Center for Translational & Computational Neuroimmunology Multiple Sclerosis Center Department of Neurology Columbia University Medical Center 630 W 168th Street P&S Box 16 New York, NY 10032. Current address: Vertex Pharmaceuticals, 50 Northern Avenue, Boston, MA 02210, USA.

## Abstract

We assembled and analyzed genetic data of 47,351 multiple sclerosis (MS) subjects and 68,284 control subjects and establish a reference map of the genetic architecture of MS that includes 200 autosomal susceptibility variants outside the major histocompatibility complex (MHC), one chromosome X variant, and 32 independent associations within the extended MHC. We used an ensemble of methods to prioritize up to 551 potentially associated MS susceptibility genes, that implicate multiple innate and adaptive pathways distributed across the cellular components of the immune system. Using expression profiles from purified human microglia, we do find enrichment for MS genes in these brain - resident immune cells. Thus, while MS is most likely initially triggered by perturbation of peripheral immune responses the functional responses of microglia and other brain cells are also altered and may have a role in targeting an autoimmune process to the central nervous system.

**One Sentence Summary:** We report a detailed genetic and genomic map of multiple sclerosis, and describe the role of putatively affected genes in the peripheral immune system and brain resident microglia.

## Introduction

Over the last decade, elements of the genetic architecture of multiple sclerosis (MS) susceptibility have gradually emerged from genome - wide and targeted studies.(*1-5*) The role of the adaptive arm of the immune system, particularly its CD4 + T cell component has become clearer, with multiple different T cell subsets being implicated.(*4*) While the T cell component plays an important role, functional and epigenomic annotation studies have begun to suggest that other elements of the immune system may be involved as well.(*6, 7*) Here, we assemble available genome-wide MS data to perform a meta-analysis followed by a systematic, comprehensive replication effort in large independent sets of subjects. This effort has yielded a detailed genome-wide genetic map that includes the first successful evaluation of the X chromosome in MS and provides a powerful platform for the creation of a detailed genomic map outlining the functional consequence of each variant and their assembly into susceptibility networks.

## Discovery and replication of genetic associations

We organized available (*1, 2, 4, 5*) and newly genotyped genome-wide data in 15 data sets, totaling 14,802 subjects with MS and 26,703 controls for our discovery study (Supplementary Methods, Supplementary Tables 1-3). After rigorous per data set quality control, we imputed all samples using the 1000 Genomes European panel resulting in an average of 8.6 million imputed single nucleotide polymorphisms (SNPs) with minor allele frequency (MAF) of at least 1% (Supplementary Methods). We then performed a meta-analysis, penalized for within-data set residual genomic inflation, to a total of 8,278,136 SNPs with data in at least two data sets (Supplementary Methods). Of these, 26,395 SNPs reached genome-wide significance (p-value < 5x10^-8^) and another 576,204 SNPs had at least nominal evidence of association (5 x 10^-8^ > p-value < 0.05). In order to identify statistically independent SNPs in the discovery set and to prioritize variants for replication, we applied a genome partitioning approach (Supplementary Methods). Briefly, we first excluded an extended region of ∼12Mb around the major histocompatibility complex (MHC) locus to scrutinize this unique region separately (see below), and we then applied an iterative method to discover statistically independent SNPs in the rest of the genome using conditional modeling. We partitioned the genome into regions by extracting ±1Mbs on either side of the most statistically significant SNP and repeating this procedure until there were no SNPs with a p-value<0.05 in the genome. Within each region we applied conditional modeling to identify statistically independent effects. As a result, we identified 1,961 non-MHC autosomal regions that included 4,842 presumably statistically independent SNPs. We refer these 4,842 prioritized SNPs as “*effects”*, assuming that these SNPs tag a true causal genetic effect. Of these, 82 effects were genome-wide significant in the discovery analysis, and another 125 had a p-value < 1 x 10^-5^5.

In order to replicate these 4,842 effects we analyzed two large-scale independent sets of data. First, we designed the MS Chip to directly replicate each of the prioritized effects (Supplementary Methods) and, after stringent quality check (Supplementary Methods and Supplementary Table 4), analyzed 20,282 MS subjects and 18,956 controls, which are organized in 9 data sets. Second, we incorporated targeted genotyping data generated using the ImmunoChip platform on an additional 12,267 MS subjects and 22,625 control subjects that had not been used in either the discovery or the MS Chip subject sets (Supplementary Table 5).(*3*) Overall, we jointly analyzed data from 47,351 MS cases and 68,284 control subjects to provide the largest and most comprehensive genetic evaluation of MS susceptibility to date.

For 4,311 of the 4,842 effects (89%) that were prioritized in the discovery analysis, we could identify at least one tagging SNP (Supplementary Methods) in the replication data. 157 regions had at least one genome-wide effect with overall 200 prioritized effects reaching a level of genome-wide significance (GW) (Figure 1). 61 of these 200 represent secondary, independent, effects that emerge from conditional modeling within a given locus (Supplementary Results and Supplementary Table 6). The odds ratios (ORs) of these genome-wide effects ranged from 1.06 to 2.06, and the allele frequencies of the respective risk allele from 2.1% to 98.4% in the European samples of the 1000 Genomes reference (mean: 51.3%, standard deviation: 24.5%;

**Figure 1.**
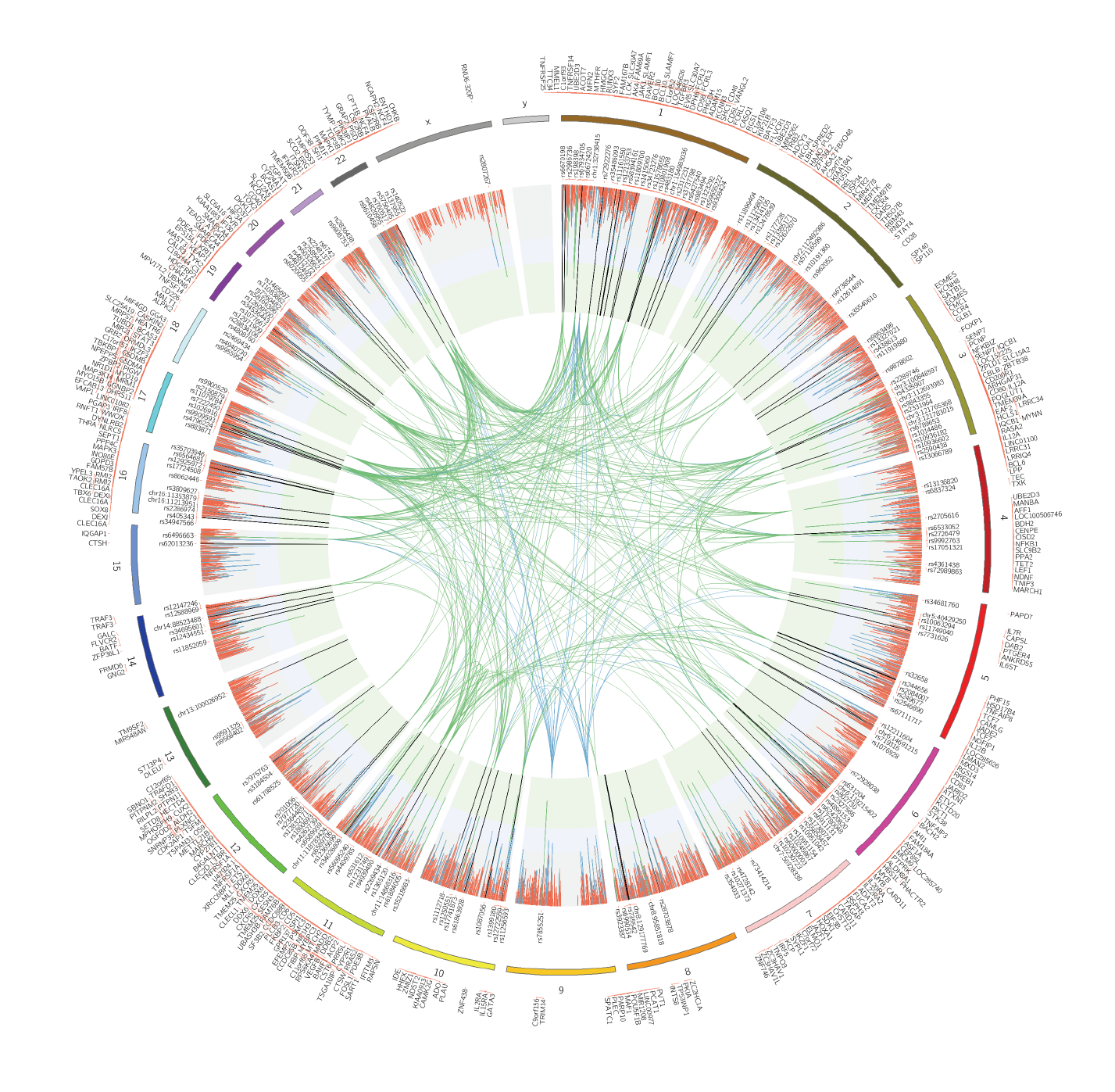
The genetic map of multiple sclerosis. The circos plot displays the 4,842 prioritized autosomal non-MHC effects and the associations in chromosome X. Joint analysis (discovery and replication) p-values are plotted as lines. The green inner layer displays genome-wide significance (p-value<5x10^-8^), the blue inner layer suggestive p-values (1x10^-5^<p-value>5x10^-8^), and the grey p-values > 1x10^-5^. Each line in the inner layers represents one effect. 200 autosomal non-MHC and one in chromosome X genome-wide effects are listed. The vertical lines in the inner layers represent one effect and the respective color displays the replication status (see main text and Online Methods): green (genome-wide), blue (potentially replicated), red (non-replicated). 551 prioritized genes are plotted on the outer surface. The inner circle space includes protein-protein interactions (PPI) between genome-wide genes (green), and genome-wide genes and potentially replicated genes (blue) that are identified as candidates using protein-protein interaction networks (see main text and Supplementary Results).

Supplementary Table 7 and Supplementary Figure 1). 19.7% of regions (31 out of 157) harbored more than one statistically independent GW effect. One of the most complex regions was the one harboring the *EVI5* gene that has been the subject of several reports with contradictory results.(*8-11*) In this locus, we identified four statistically independent genome-wide effects, three of which were found under the same association peak (Figure 2A), illustrating how our approach and the large sample size clarifies associations described in smaller studies and can facilitate functional follow-up of complex loci.

**Figure 2.**
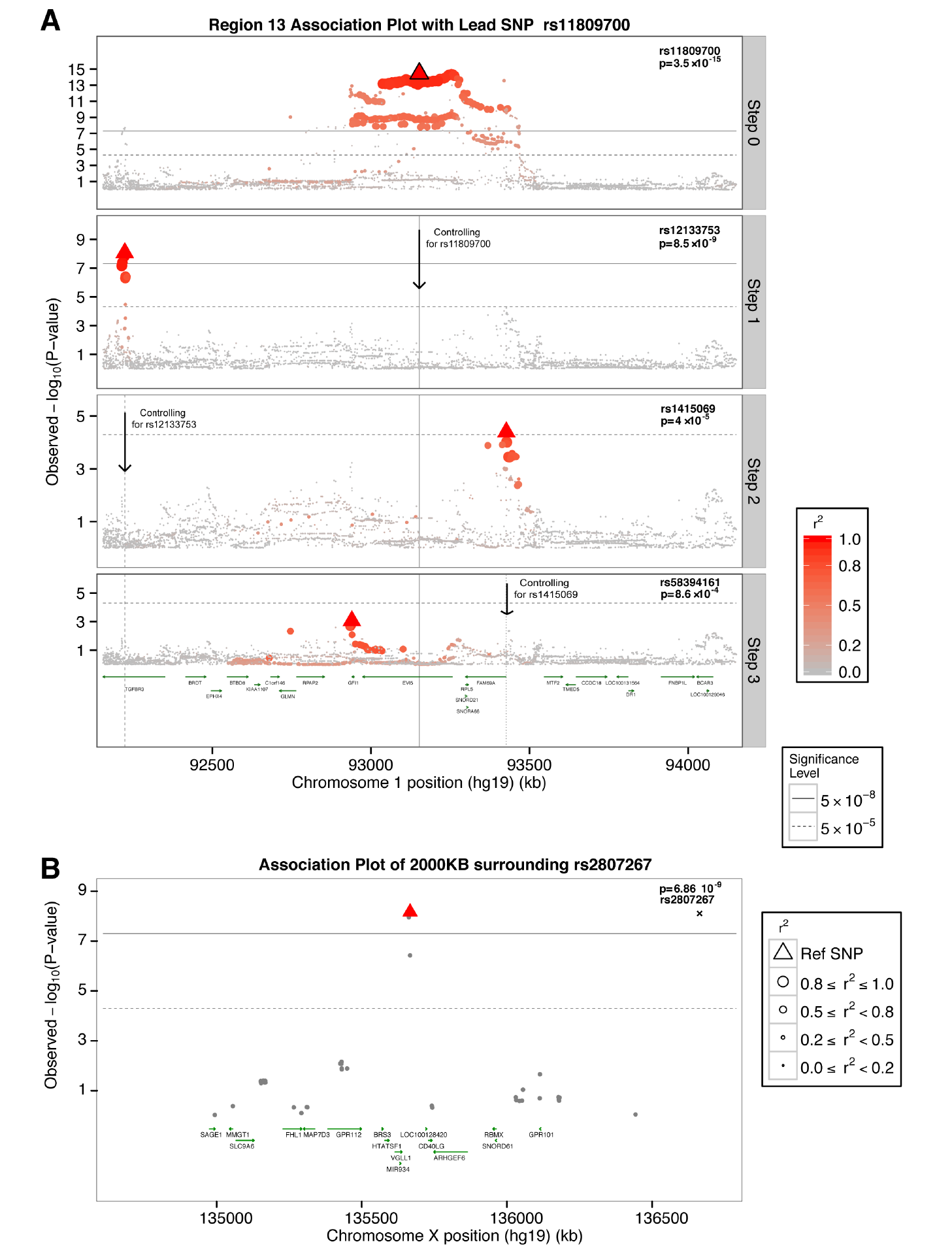
Multiple independent effects in the *EVI5* locus and chromosome X associations. **A)** Regional association plot of the *EVI5* locus. Discovery p-values are displayed. The layer tagged “Marginal” plots the associations of the marginal analysis, with most statistically significant SNP being rs11809700 (OR_T_ = 1.16; p-value = 3.51x10^-15^). The “Step 1” plots the associations conditioning on rs11809700; rs12133753 is the most statistically significant SNP (OR_C_ = 1.14; p-value = 8.53x10^-09^). “Step 2” plots the results conditioning on rs11809700 and rs12133753, with rs1415069 displaying the lowest p-value (OR_G_ = 1.10; p-value = 4.01x10^-5^). Finally, “Step 3” plots the associations conditioning on rs11809700, rs12133753, and rs1415069, identifying rs58394161 as the most-statistically significant SNP (OR_C_ = 1.10; p-value = 8.63x10^-4^4). All 4 SNPs reached genome-wide significance in the respective joint, discovery plus replication, analyses (Supplementary Table 6). Each of the independent 4 SNPs, i.e. lead SNPs, are highlighted using a triangle in the respective layer. **B**) Regional association plot for the genome-wide chromosome X variant. Joint analysis p-values are displayed. Linkage disequilibrium, in terms of r^2^ based on the 1000 Genomes European panel, is indicated using a combination of color grade and symbol size (see legend for details). All positions are in human genome 19.

We also performed a joint analysis of available data on sex chromosome variants (Supplementary Methods) and we identified rs2807267 as genome-wide significant (OR_T_ = 1.07, p-value = 6.86 x 10^-9^; Supplementary Tables 8-9). This variant lies within an enhancer peak specific for T cells and is 948bps downstream of the RNA U6 small nuclear 320 pseudogene (*RNU6-320P*), a component of the U6 snRNP (small nuclear ribonucleoprotein) that is part of the spliceosome and is responsible for the splicing of introns from pre-mRNA (*12*) (Figure 2B). The nearest gene is *VGLL1* (27,486bps upstream) that has been proposed to be a co-activator of mammalian transcription factors.(*13*) No variant in the Y chromosome had a p-value lower than 0.05 in either the discovery or replication sets.

The MHC was the first MS susceptibility locus to be identified, and prior studies have found that it harbors multiple independent susceptibility variants, including interactions within the class II HLA genes.(*14, 15*) We undertook a detailed modeling of this region to account for its long-range linkage disequilibrium and allelic heterogeneity using SNP data as well as imputed classical alleles and amino acids of the human leukocyte antigen (HLA) genes in the assembled data. We confirm prior MHC susceptibility variants (including a non-classical HLA effect located in the *TNFA / LST1* long haplotype) and we extend the association map to uncover a total of 31 statistically independent effects at the genome-wide level within the MHC (Figure 3, Supplementary Table 10). An interesting finding is that several HLA and nearby non-HLA genes have several independent effects that can now be identified due to our large sample, e.g. the *HLA-DRB1* locus has six statistically independent effects. Another exciting finding involves *HLA-B* that also appears to harbor 6 independent effects on MS susceptibility. The role of the non-classical HLA and non-HLA genome in the MHC is also highlighted. One third (9 out of 30) of the identified variants lie within either intergenic regions or in a long-range haplotype that contains several non-classical HLA and other non-HLA genes.(*15*) Recently, we reported an interaction between HLA-DRB1*15:01 and HLA-DQA1*01:01 by analyzing imputed HLA alleles.(*14*) Here we reinforce this analysis by analyzing SNPs, HLA alleles, and respective amino acids. We replicate the presence of interactions among class II alleles but note that the second interaction term, besides HLA-DRB1*15:01, can vary depending on the other independent variants that are included in the model. First, we found that there are interaction models of HLA-DRB1*15:01 with other variants in MHC that explain better the data than our previously reported HLA-DRB1*15:01 / HLA-DQA1*01:01 interaction term (Supplementary Figure 2). Second, we observe that there is a group of *HLA*DQB1* and *HLA*DQA1* SNPs, alleles, and amino acids that consistently rank amongst the best models with HLA-DRB1*15:01 interaction terms (Supplementary Figure 3). This group of HLA-DRB1*15:01-interacting variants is consistently identified regardless of the marginal effects of other statistically independent variants that are added in the model, implying that these interaction terms capture a different subset of phenotypic variance and can be explored after the identification of the marginal effects. Finally, we performed a sensitivity analysis by including interaction terms of HLA-DRB1*15:01 in each step and selecting the model with the lowest Bayesian information criterion (BIC), instead of testing only the marginal results of the variants as we did in the main analysis. This sensitivity analysis also resulted in 32 statistically independent effects with a genome-wide significant p-value (Supplementary Table 11), of which one third (9 out of 32) were not effects in classical HLA genes. The main differences between the results of the two approaches were the inclusion of interaction of HLA-DRB1*15:01 and rs1049058 in step 3 and the stronger association of *HLA*DPB1/2* effects over *HLA*DRB1* effects in the sensitivity model (Supplementary Tables 11-12 and Supplementary Figure 3). Thus, overall, our MHC results are not strongly affected by the analytic model that we have selected.

**Figure 3.**
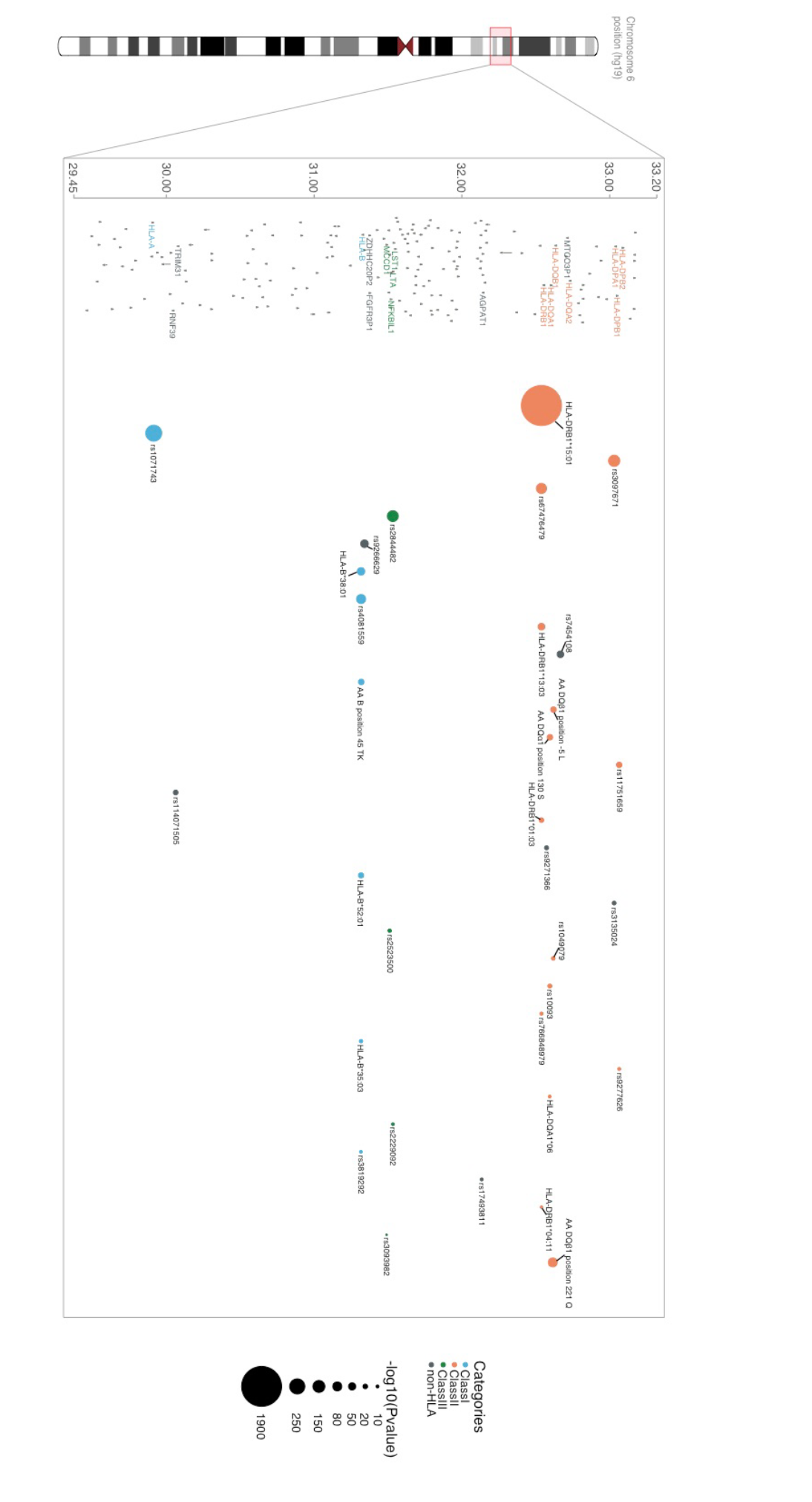
Independent associations in the major histocompatibility locus. Regional association plot in the MHC locus. Only genome-wide statistically independent effects are listed. The order of variants in the X-axis represents the order these were identified. The size of the circle represents different values of –log10(p-value). Different colors are used to depict class I, II, III, and non-HLA effects. Y-axis displays million base pairs.

## Characterization of non-genome wide effects

The commonly used threshold of genome-wide significance (p-value = 5 x 10^-8^) has played an important role in making human genetic study results robust; however, several studies have demonstrated that non-genome-wide effects explain an important proportion of the effect of genetic variation on disease susceptibility. (*16, 17*) More importantly, several such effects are eventually identified as genome-wide significant, given enough sample size and true effects.(*3*) Thus, we also evaluated the non-genome-wide effects that were selected for replication, have available replication data (n = 4,116), but do not meet a standard threshold of genome-wide significance (p<5x10^-8^). Specifically, we decided to stratify these 4,116 effects into 2 main categories (see details in Supplementary Methods): (1) suggestive effects (S, n = 416), and (2) non-replicated effects (NR, n = 3,694). We used these categories in downstream analyses to further characterize the prioritized effects from the discovery study in terms of potential to eventually be replicated. We also included a third category: effects for which there were no data for replication in any of the replication sets (no data, ND, n = 532). Furthermore, to add granularity in each category, we sub-stratified the suggestive effects into 2 groups: (1a) strongly suggestive (5 x10^-8^ > p-value <1x10^-^5; sS, n = 118) and (1b) underpowered suggestive (unS, n = 299). Of these two categories of suggestive effects, the ones in the sS category have a high probability of reaching genome-wide significance as we increase our sample size in future studies (Supplementary Results and Supplementary Table 13).

## Heritability explained

To estimate the extent to which we have characterized the genetic architecture of MS susceptibility with our 200 genome-wide non-MHC autosomal MS effects, we calculated the narrow-sense heritability captured by common variation (*h2g*), i.e. the ratio of additive genetic variance to the total phenotypic variance (Supplementary Methods).(*16, 18*) Only the 15 strata of data from the discovery set had true genome-wide coverage, and hence we used these 14,802 MS subjects and 26,703 controls for the heritability analyses. The overall heritability estimate for MS susceptibility in the discovery set of subjects was 19.2% (95%CI: 18.5-19.8%). Heritability partitioning using minor allele frequency or p-value thresholds has led to significant insights in previous studies,(*19*) and we therefore applied a similar partitioning approach but in a fashion that took into consideration the study design and the existence of replication information from the 2 large-scale replication cohorts. First, we partitioned the autosomal genome into 3 components: i) the super extended MHC (SE MHC, see above), ii) a component with the 1,961 regions prioritized for replication (Regions), and iii) the rest of the genome that had p-value>0.05 in the discovery study (Non-associated regions). Then, we estimated the *h2g* that can be attributed to each component as a proportion of the overall narrow-sense heritability observed. The SE MHC explained 21.4% of the *h2g*, with the remaining 78.6% being captured by the second component (Figure 4A). Then, we further partitioned the non-MHC component into one that captured all 4,842 statistically independent effects (Prioritized for replication), which explained the vast majority of the overall estimated heritability: 68.3%. The “Non-prioritized” SNPs in the 1,961 regions explained 11.6% of the heritability, which suggests that there may be residual LD with prioritized effects or true effects that have not yet been identified (Figure 4B).

**Figure 4.**
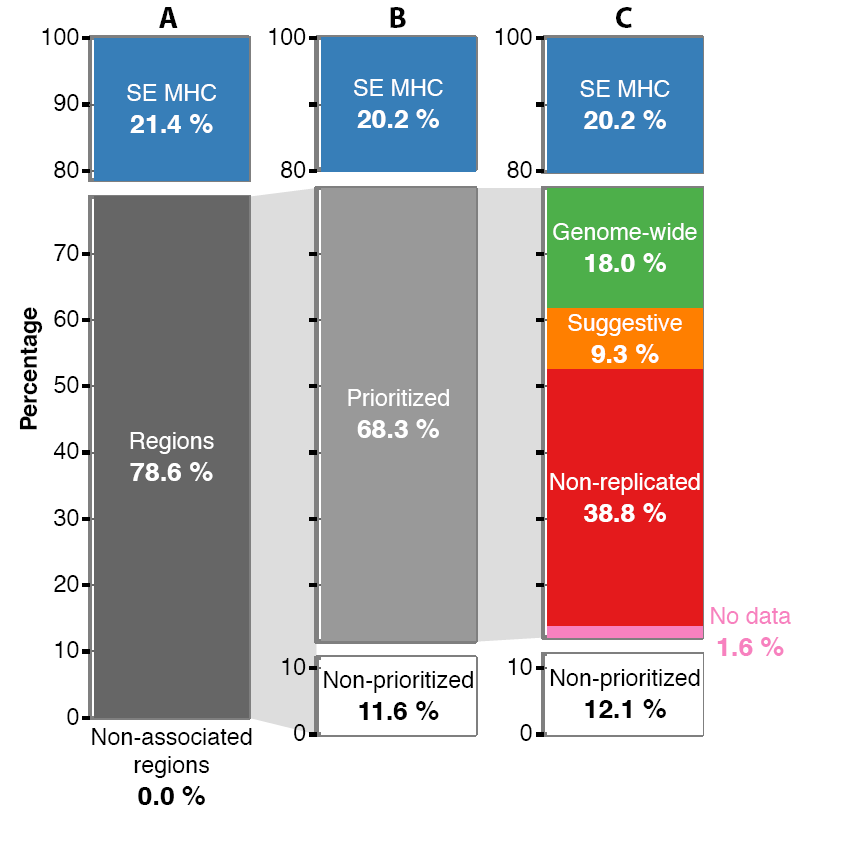
Heritability partitioning. Proportion of the overall narrow-sense heritability under the liability model (∼19.2%) explained by different genetic components. (A) The overall heritability is partitioned in the super extended MHC (SE MHC), the 1,962 Regions that include all SNPs with p-value<0.05 (Regions), and the rest of genome with p-values>0.05 (Non-associated regions). (B) The Regions are further partitioned to the seemingly statistically independent effects (Prioritized) and the residual (Non-prioritized). (C) The Prioritized component is partitioned based on the replication knowledge to genome-wide effects (GW), suggestive (S), non-replicated (ND), and no data (ND). The lines connecting the pie charts depict the component that is partitioned. All values are estimated using the discovery data-sets (n = 4,802 cases and 26,703 controls).

We then used the replication-based categories described above to further partition the “Prioritized” heritability component, namely “GW”, “S”, “NR”, “ND” (Figure 4C). The genome-wide effects (GW) captured 18.3% of the overall heritability. Thus, along with the contribution of the SE MHC (20.2% in the same model), we can now explain ∼39% of the genetic predisposition to MS with the validated susceptibility alleles. This can be extended to ∼48% if we include the suggestive (S) effects (9.0%). Interestingly the non-replicated (NR) effects captured 38.8% of the heritability, which could imply that some of these effects might be falsely non-replicated, i.e. that these are true effects that need further data to emerge robustly or that their effect may be true and present in only a subset of the data. However, few of the 3,694 NR effects would fall in either of the above two cases; the vast majority of these effects are likely to be false positive results.

## Functional implications of the MS loci, enriched pathways and gene-sets

Next, we began to annotate the MS effects. To prioritize the cell types or tissues in which the 200 non-MHC autosomal effects may exert their effect, we used two different approaches: one that leverages atlases of gene expression patterns and another that uses a catalog of epigenomic features such as DNase hypersensitivity sites (DHSs) (Supplementary Methods).(*7, 20-22*) Significant enrichment for MS susceptibility loci was apparent in many different immune cell types and tissues, whereas there was an absence of enrichment in tissue-level central nervous system (CNS) profiles (Figure 5). An important finding is that the enrichment is observed not only in immune cells that have long been studied in MS, e.g. T cells, but also in B cells whose role has emerged more recently.(*23*) Furthermore, while the adaptive immune system has been proposed to play a predominant role in MS onset,(*24*) we now demonstrate that many elements of innate immunity, such as natural killer (NK) cells and dendritic cells also display strong enrichment for MS susceptibility genes. Interestingly, at the tissue level, the role of the thymus is also highlighted, possibly suggesting the role of genetic variation in thymic selection of autoreactive T cells in MS.(*25*) Public tissue-level CNS data – which are derived from a complex mixture of cell types-do not show an excess of MS susceptibility variants in annotation analyses. However, since MS is a disease of the CNS, we extended the annotation analyses by analyzing new data generated from human iPSC-derived neurons as well as from purified primary human astrocytes and microglia. As seen in Figure 6, enrichment for MS genes is seen in human microglia (p = 5 x 10^-14^) but not in astrocytes or neurons, suggesting that the resident immune cells of the brain may also play a role in MS susceptibility

**Figure 5.**
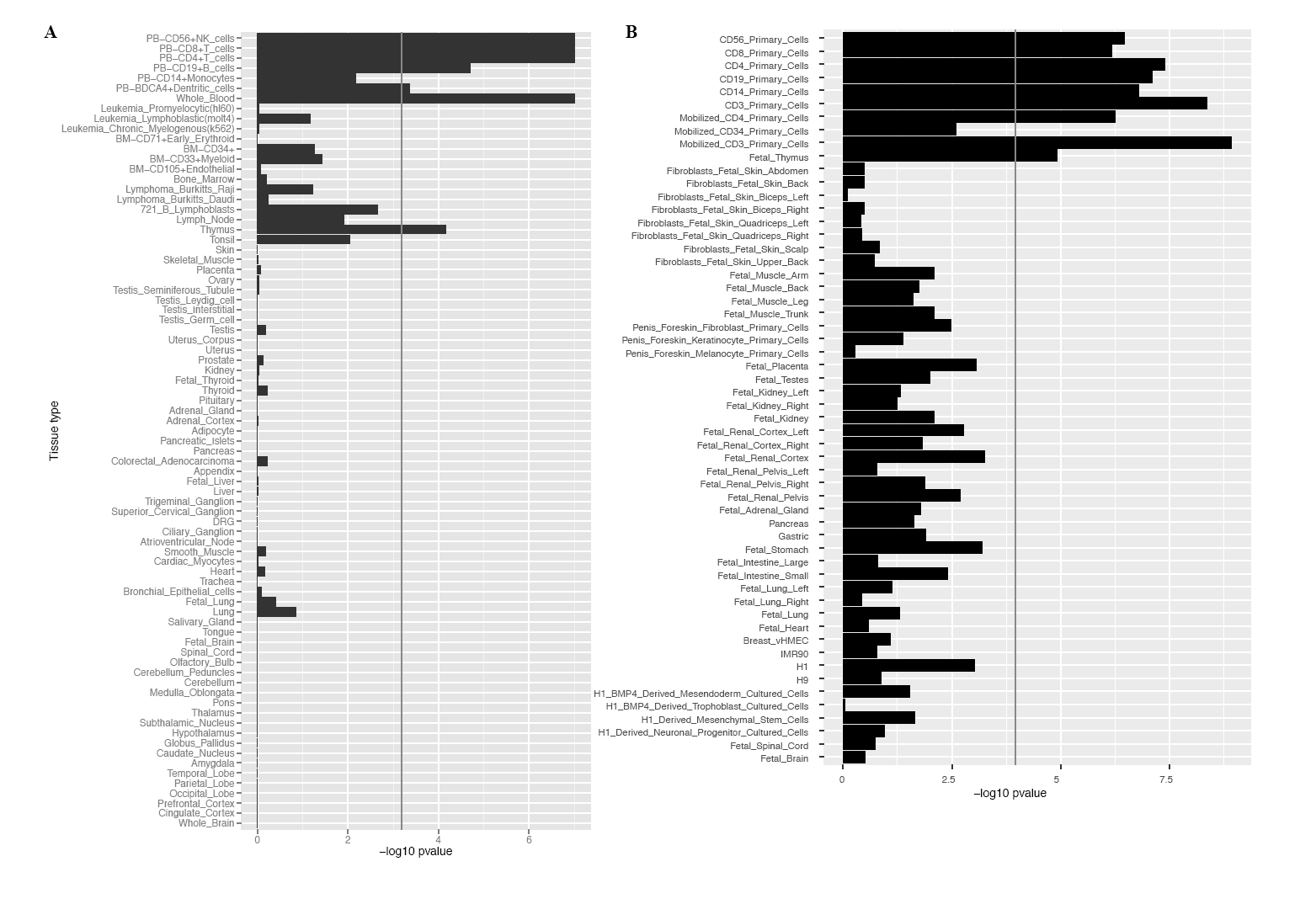
Tissue and cell type enrichment analyses. (A) Gene Atlas tissues and cell types gene expression enrichment. (B) DNA hypersensitivity sites (DHS) enrichment for tissues and cell types from the NIH Epigenetic Roadmap. Rows are sorted from immune cells/tissues to central nervous system related ones. Both X axes display –log10 of Benjamini & Hochberg p-values (false discovery rate).

**Figure 6.**
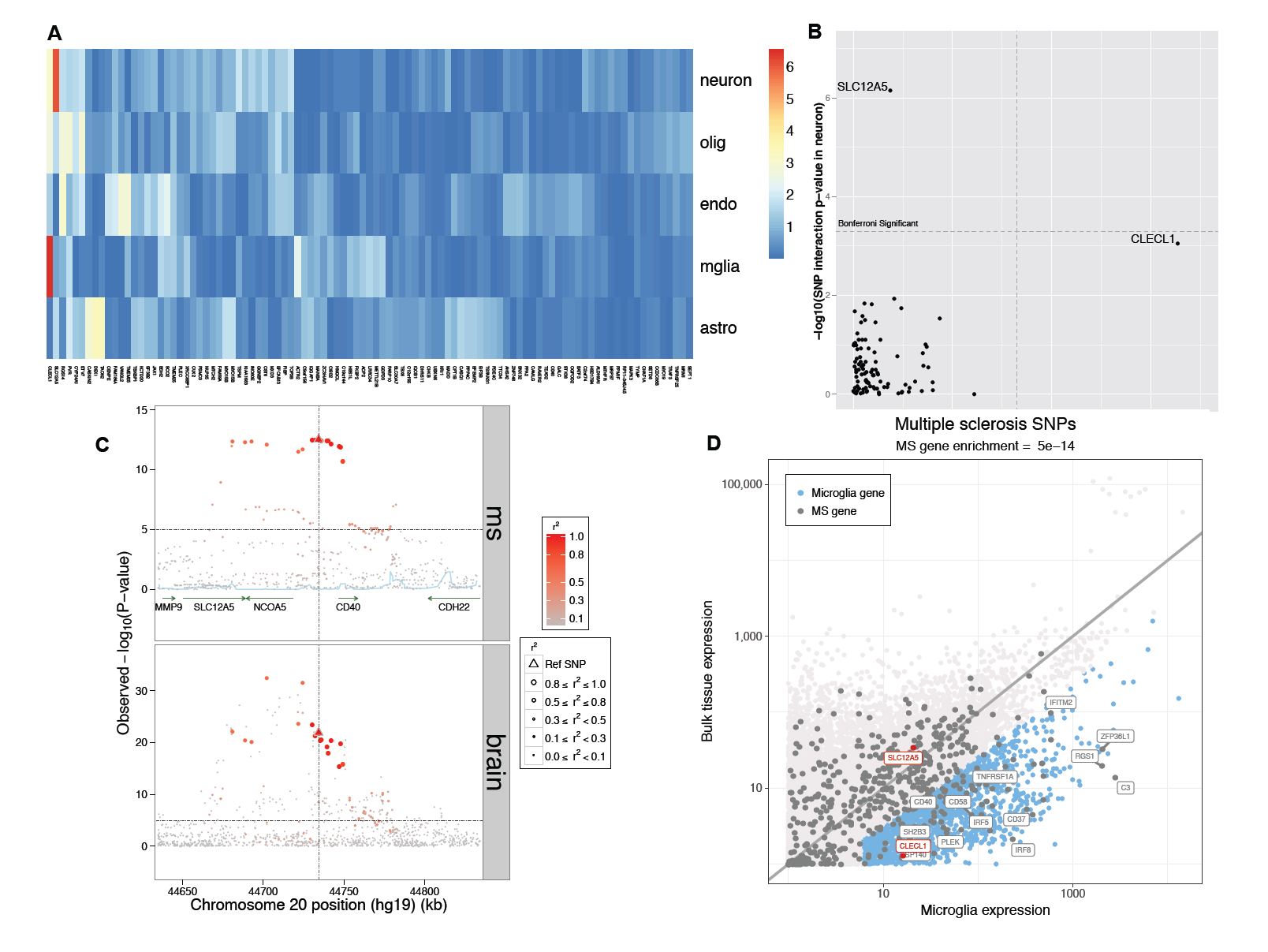
Dissection of cortical RNAseq data. In (A), we present a heatmap of the results of our analysis assessing whether a cortical eQTL is likely to come from one of the component cell types of the cortex: neurons, oligodendrocytes, endothelial cells, microglia and astrocytes (in rows). Each column presents results for one of the MS brain eQTLs. The color scheme relates to the p-value of the interaction term, with red denoting a more extreme result. (B) We present the same results in a different form, comparing results of assessing for interaction with neuronal proportion (y axis) and microglial proportion (x-axis): the *SLC12A5* eQTL is significantly stronger when accounting for neuronal proportion, and *CLECL1* is significantly stronger when accounting for microglia. The Bonferroni-corrected threshold of significance is highlighted by the dashed line. (C) Locus view of the *SLC12A5/CD40* locus, illustrating the distribution of MS susceptibility and the *SLC12A5* brain eQTL in a segment of chromosome 20 (x axis); the y axis presents the p-value of association with MS susceptibility (top panel) or *SLC12A5* RNA expression (bottom panel). The lead MS SNP is denoted by a triangle, other SNPs are circles, with the intensity of the red color denoting the strength of LD with the lead MS SNP in both panels. (D) Here we plot the level of expression, transcriptome-wide, for each measured gene in our cortical RNAseq dataset (n = 455)(y-axis) and purified human microglia (n = 10)(x-axis) from the same cortical region. In blue, we highlight those genes with > 4 fold increased expression in microglia relative to bulk cortical tissue and are expressed at a reasonable level in microglia. Each dot is one gene. Gray dots denote the 551 putative MS genes from our integrated analysis. *SLC12A5* and *CLECL1* are highlighted in red; in blue, we highlight a selected subset of the MS genes – many of them well-validated – which are enriched in microglia. For clarity, we did not include all of the MS genes that fall in this category.

We repeated the enrichment analyses for the “S” and “NR” effects aiming to test whether these have a similar enrichment pattern with the 200 “GW” effects. The “S” effects exhibited a pattern of enrichment that is similar to the “GW” effects, with only B cell expression reaching a threshold of statistical significance (Supplementary Figure 4). This provides additional circumstantial evidence that this category of variants may harbor true causal associations. On the other hand, the “NR” enrichment results seem to follow a rather random pattern, suggesting that most of these effects are indeed not truly MS-related (Supplementary Figure 4).

The strong enrichment of the GW effects in immune cell types motivated us to prioritize candidate MS susceptibility genes by identifying those susceptibility variants, which affect RNA expression of nearby genes (*cis* expression quantitative trait loci effect, *cis -* eQTL) (±500Kbps around the effect SNP; Supplementary Methods). Thus, we interrogated the potential function of MS susceptibility variants in naive CD4 + T cells and monocytes from 415 healthy subjects as well as peripheral blood mononuclear cells (PBMCs) from 225 remitting relapsing MS subjects. Thirty-six out of the 200 GW MS effects (18%) had at least one tagging SNP (r2> = 0.5) that altered the expression of 46 genes (FDR<5%) in CD4 + naïve T cells (Supplementary Table 14), and 36 MS effects (18%; 10 common with the CD4 + naïve T cells) influenced the expression of 48 genes in monocytes (11 genes in common with T cells). In MS PBMC, 30% of the GW effects (60 out of the 200) were cis-eQTLs for 92 genes in the PBMC MS samples, with several loci being shared with those found in healthy T cells and monocytes (26 effects and 27 genes in T cells, and 21 effects and 24 genes in monocytes, respectively; Supplementary Table 14).

Since MS is a disease of the CNS, we also investigated a large collection of dorsolateral prefrontal cortex RNA sequencing profiles from two longitudinal cohort studies of aging (n = 455), which recruit cognitively non-impaired individuals (Supplementary Methods). This cortical sample provides a tissue-level profile derived from a complex mixture of neurons, astrocytes, and other parenchymal cells such as microglia and occasional peripheral immune cells. In these data, we found that 66 of the GW MS effects (33% of the 200 effects) were *cis*-eQTLs for 104 genes. Over this CNS and the three immune sets of data, 104 GW effects were *cis*-eQTLs for 203 unique genes (n = 211 *cis*-eQTLs), with several appearing to be seemingly specific for one of the cell/tissue type (Supplementary Table 14). Specifically, 21.2**%** (45 out of 212 *cis*-eQTLs) of these cortical cis-eQTLs displayed no evidence of association (p-value>0.05 with any SNP with r^2^>0.1) in the immune cell/tissue results and are less likely to be immune-related (Supplementary Table 15).

To further explore the challenging and critical question of whether some of the MS variants have an effect that is primarily exerted through a non-immune cell, we performed a secondary analysis of our cortical RNAseq data in which we attempted to ascribe a brain cis-eQTL to a particular cell type. Specifically, we assessed our tissue-level profile and adjusted each cis-eQTL analysis for the proportion of neurons, astrocytes, microglia, and oligodendrocytes estimated to be present in the tissue: the hypothesis was that the effect of a SNP with a cell type-specific cis-eQTL would be stronger if we adjusted for the proportion of the target cell type (Figure 6). As anticipated, almost all of the MS variants present in cortex remain ambiguous: it is likely that many of them influence gene function in multiple immune and non-immune cell types. However, the *SLC12A5* locus is different: here, the effect of the SNP is significantly stronger when we account for the proportion of neurons (Figure 6A and 6B), and the *CLECL1* locus emerges when we account for the proportion of microglia. *SLC12A5* is a potassium / chloride transporter that is known to be expressed in neurons, and a rare variant in *SLC12A5* causes a form of pediatric epilepsy (*26, 27*). While this MS locus may therefore appear to be a good candidate to have a primarily neuronal effect, further evaluation finds that this MS susceptibility haplotype also harbors susceptibility to rheumatoid arthritis (*28*) and a cis-eQTL in B cells for the *CD40* gene (*29*). Thus, the same haplotype harbors different functional effects in very different contexts, illustrating the challenge in dissecting the functional consequences of autoimmune variants in immune as opposed to the tissue targeted in autoimmune disease. On the other hand, *CLECL1* represents a simpler case of a known susceptibility effect that has previously been linked to altered *CLECL1* RNA expression in monocytes (*24, 30*); its enrichment in microglial cells, which share many molecular pathways with other myeloid cells, is more straightforward to understand. *CLECL1* is expressed at low level in our cortical RNAseq profiles because microglia represent just a small fraction of cells at the cortical tissue level, and its expression level is 20-fold greater when we compare its level of expression in purified human cortical microglia to the bulk cortical tissue (Figure 6). *CLECL1* therefore suggests a potential role of microglia in MS susceptibility, which is under-estimated in bulk tissue profiles that are available in epigenomic and transcriptomic reference data. Overall, many genes that are eQTL targets of MS variants in the human cortex are most likely to affect multiple cell types. These brain eQTL results and the enrichment found in analyses of our purified human microglia data therefore highlight the need for more targeted, cell-type specific data for the CNS to adequately determine the extent of its role in MS susceptibility.

These eQTL studies begin to transition our genetic map into a resource outlining the likely MS susceptibility gene(s) in a locus and the potential functional consequences of certain MS variants. To assemble these single-locus results into a higher-order perspective of MS susceptibility, we turned to pathway analyses to evaluate how the extended list of genome-wide effects provides new insights into the pathophysiology of the disease. Acknowledging that there is no available method to identify all causal genes following GWAS discoveries, we prioritized genes for pathway analyses while allowing several different hypotheses for mechanisms of actions (Supplementary Methods). In brief, we prioritized genes that: (i) were *cis*-eQTLs in any of the eQTL data sets outlined above, (ii) had at least one exonic variant at r^2^> = 0.1 with any of the 200 effects**, (**iii) had high score of regulatory potential using a cell specific network approach, (iv) had a similar co-expression pattern as identified using DEPICT.(*31*) Sensitivity analyses were performed including different combinations of the above categories, and including genes with intronic variants at r^2^> = 0.5 with any of the 200 effects (Supplementary Methods and Supplementary Results). Overall, we prioritized 551 candidate MS genes (Supplementary Table 16; Supplementary Table 17 for sensitivity analyses) to test for statistical enrichment of known pathways. Approximately 39.6% (142 out of 358) of the Ingenuity Pathway Analysis (IPA) canonical pathways,(*32*) that had overlap with at least one of the identified genes, were enriched for MS genes at an FDR<5% (Supplementary Table 18). Sensitivity analyses including different criteria to prioritize genes revealed a similar pattern of pathway enrichment (Supplementary Results and Supplementary Table 19). Interestingly, the extensive list of susceptibility genes, that more than doubles the previous knowledge in MS, captures processes of development, maturation, and terminal differentiation of several immune cells that potentially interact to predispose to MS. In particular, the role of B cells, dendritic cells and natural killer cells has emerged more clearly, broadening the prior narrative of T cell dysregulation that emerged from earlier studies.(*4*) Given the over-representation of immune pathways in these databases, ambiguity remains as to where some variants may have their effect: neurons and particularly astrocytes repurpose the component genes of many “immune” signaling pathways, such as the ciliary neurotrophic factor (CNTF), nerve growth factor (NGF), and neuregulin signaling pathways that are highly significant in our analysis (Supplementary Table 18). These results – along with the results relating to microglia – emphasize the need for further dissection of these pathways in specific cell types to resolve where a variant is exerting its effect; it is possible that multiple, different cell types could be involved in disease since they all experience the effect of the variant.

Pathway and gene-set enrichment analyses can only identify statistically significant connections of genes in already reported, and in some cases validated, mechanisms of action. However, the function of many genes is yet to be uncovered and, even for well-studied genes, the full repertoire of possible mechanisms is incomplete. To complement the pathway analysis approach and to explore the connectivity of our prioritized GW genes, we performed a protein-protein interaction (PPI) analysis using GeNets (Supplementary Methods).(*33*) About a third of the 551 prioritized genes (n = 190; 34.5%) were connected (p-value = 0.052) and these could be organized into 13 communities, i.e. sub-networks with higher connectivity (p-value: < 0.002; Supplementary Table 20). This compares to 9 communities that could be identified by the previously reported MS susceptibility list (81 connected genes out of 307; Supplementary Table 21).(*3*) Next, we leveraged GeNets to predict candidate genes based on network connectivity and pathway membership similarity and test whether our previous known MS susceptibility list could have predicted any of the genes prioritized by the newly identified effects. Of the 244 genes prioritized by novel findings (out of the 551 overall prioritized genes) only five could be predicted given the old results (out of 70 candidates; Supplementary Figure 5 and Supplementary Table 22). In a similar fashion we estimated that the list of 551 prioritized genes could predict 102 new candidate genes, four of which can be prioritized since they are in the list of suggestive effects. (Figure 1; Supplementary Figure 6 and Supplementary Table 23).

## Discussion

This detailed genetic map of MS is a powerful substrate for annotation and functional studies and provides a new level of understanding for the molecular events that contribute to MS susceptibility. It is clear that these events are widely distributed across the many different cellular components of both the innate and adaptive arms of the immune system: every major immune cell type is enriched for MS susceptibility genes. An important caveat is that many of the implicated molecular pathways, such as response to TNFa and type I interferons, are repurposed in many different cell types, leading to an important ambiguity: is risk of disease driven by altered function of only one of the implicated cell types or are all of them contributing to susceptibility equally? This question highlights the important issue of the context in which these variants are exerting their effects. We have been thorough in our evaluation of available reference epigenomic data, but many different cell types and cell states remain to be characterized and could alter our summary. Further, inter-individual variability has not been established in such reference data that are typically produced from one or a handful of individuals; thus, this issue is better evaluated in the eQTL data where we have examined a range of samples and states in large numbers of subjects. Overall, while we have identified putative functional consequences for the identified MS variants, the functional consequence of most of these MS variants requires further investigation.

Even where a function is reported, further work is needed to demonstrate that the effect is the causal functional change. This is particularly true of the role of the CNS in MS susceptibility: we mostly have data at the level of the human cortex, a complex tissue with many different cell types, including resident microglia and a small number of infiltrating macrophage and lymphocytes. MS variants clearly influence gene expression in this tissue, and we must now: (1) resolve the implicated cell types and whether pathways shared with immune cells are having their MS susceptibility effect in the periphery or in the brain and; (2) more deeply identify additional functional consequences that may be present in only a subset of cells, such as microglia or activated astrocytes, that are obscured in the cortical tissue level profile. A handful of loci are intriguing in that they alter gene expression in the human cortex but not in the sampled immune cells; these MS susceptibility variants deserve close examination to resolve the important question of the extent to which the CNS is involved in disease onset. Thus, our study suggests that while MS is a disease whose origin may lie primarily within the peripheral immune compartment where dysregulation of all branches of the immune system leads to organ specific autoimmunity, there is subset of loci with a key role in directing the tissue specific autoimmune response. This is similar to our previous examination of ulcerative colitis, where we observed enrichment of genetic variants mapping to colon tissue.(*6*) This view is consistent with our understanding of the mechanism of important MS therapies such as natalizumab and fingolimod that sequester pathogenic immune cell populations in the peripheral circulation to prevent episodes of acute CNS inflammation. It also has important implications as we begin to consider prevention strategies to block the onset of the disease by early targeting peripheral immune cells.

An important step forward in MS genetics, for a disease with a 3:1 preponderance of women being affected, is robust evidence for a susceptibility locus on the X chromosome. The function of this locus remains to be defined in future studies. Nonetheless, it provides a key first step for a genetic component to the role of sex, which is the risk factor of largest effect in MS susceptibility.(*34*) This result also highlights the need for additional, dedicated genetic studies of the X chromosome in MS as existing data have not been fully leveraged. (*35*)

This genomic map of MS – the genetic map and its integrated functional annotation-is a foundation on which the next generation of projects will be developed. Beyond the characterization of the molecular events that trigger MS, this map will also inform the development of primary prevention strategies since we can leverage this information to identify the subset of individuals who are at greatest risk of developing MS. While insufficient by itself, an MS Genetic Risk Score has a role to play in guiding the management of the population of individuals “at risk” of MS (such as family members) when deployed in combination with other measures of risk and biomarkers that capture intermediate phenotypes along the trajectory from health to disease.(*36*) We thus report an important milestone in the investigation of MS and share a roadmap for future work: the establishment of a map with which to guide the development of the next generation of studies with high-dimensional molecular data to explore both the initial steps of immune dysregulation across both the adaptive and innate arms of the immune system and secondly the translation of this auto-immune process to the CNS where it triggers a neurodegenerative cascade.

## Acknowledgments

This investigation was supported in part by a Postdoctoral Fellowship from the National Multiple Sclerosis Society (FG 1938-A-1) and a Career Independence Award from the National Multiple Sclerosis Society (TA 3056-A-2) to Nikolaos A. Patsopoulos. Nikolaos A. Patsopoulos has been supported by Harvard NeuroDiscovery Center and an Intel Parallel Computing Center award. Swedish Medical Research Council; Swedish Research Council for Health, Working Life and Welfare, Knut and Alice Wallenberg Foundation, AFA insurance, Swedish Brain Foundation, the Swedish Association for Persons with Neurological Disabilities. This study makes use of data generated as part of the Wellcome Trust Case Control Consortium 2 project (085475/B/08/Z and 085475/Z/08/Z), including data from the British 1958 Birth Cohort DNA collection (funded by the Medical Research Council grant G0000934 and the Wellcome Trust grant 068545/Z/02) and the UK National Blood Service controls (funded by the Wellcome Trust). The study was supported by the Cambridge NIHR Biomedical Research Centre, UK Medical Research Council (G1100125) and the UK MS society (861/07). NIH/NINDS: R01 NS049477, NIH/NIAID: R01 AI059829, NIH/NIEHS: R01 ES0495103. Research Council of Norway grant 196776 and 240102. NINDS/NIH R01NS088155. Oslo MS association and the Norwegian MS Registry and Biobank and the Norwegian Bone Marrow Registry. Research Council KU Leuven, Research Foundation Flanders. AFM, AFM-Généthon, CIC, ARSEP, ANR-10-INBS-01 and ANR-10-IAIHU-06. Research Council KU Leuven, Research Foundation Flanders. Inserm ATIP-Avenir Fellowship and Connect-Talents Award. German Ministry for Education and Research, German Competence Network MS (BMBF KKNMS). Dutch MS Research Foundation. TwinsUK is funded by the Wellcome Trust, Medical Research Council, European Union, the National Institute for Health Research (NIHR)-funded BioResource, Clinical Research Facility and Biomedical Research Centre based at Guy’s and St Thomas’ NHS Foundation Trust in partnership with King’s College London. We thank the volunteers from the Oxford Biobank(www.oxfordbiobank.org.uk) and the Oxford NIHR Bioresource for their participation. The recall process was supported by the National Institute for Health Research (NIHR) Oxford Biomedical Research Centre Programme. The views expressed are those of the author(s) and not necessarily those of the NHS, the NIHR or the Department of Health. German Ministry for Education and Research, German Competence Network MS (BMBF KKNMS). Italian Foundation of Multiple Sclerosis (FISM). NMSS (RG 4680A1/1). German Ministry for Education and Research, German Competence Network MS (BMBF KKNMS). Lundbeck Foundation and Benzon Foundation for support (THP). This research was supported by grants from the Danish Multiple Sclerosis Society, the Danish Council for Strategic Research [grant number 2142-08-0039], Novartis, Biogen Biogen (Denmark) A/S, and the Sofus Carl Emil Friis og Hustru Olga Doris Friis Foundation and the Foundation for Research in Neurology. No conflicts of interest are reported.

The genome wide associations studies used in the discovery phase are available in the following repositories: i) dbGAP: phs000275.v1.p1, phs000139.v1.p1, phs000294.v1.p1, and phs000171.v1.p1, and ii) European Genome-phenome Archive database: EGAD00000000120, EGAD00000000022, and EGAD00000000021. The ANZGENE GWAS data is available via request to the ANZGENE Consortium. A direct request can be made via MSRA.org.au. The Rotterdam GWAS data can be requested via e-mailing. The Berkeley data set can be requested via Kaiser Permanente. The MS Chip and ImmunoChip data are under submission in dbGAP. The ImmVar data are available in GEO: GSE56035. The MS PBMC data are available in GEO: GSE16214. Human Gene Atlas: http://snpsea.readthedocs.io/en/latest/data.html#geneatlas2004-gct-gz. ImmGen: http://snpsea.readthedocs.io/en/latest/data.html#immgen2012-gct-gz. The brain related data are available in Synapse: https://www.synapse.org/#!Synapse:syn2580853/wiki/409844.

## Supplementary Materials

The following supplementary materials accompany the paper:

1) Supplementary Methods, Results and Figures

2) Supplementary Tables in excel format

## Membership of Wellcome Trust Case Control Consortium 2 (WTCCC2)

### Management Committee

Peter Donnelly (Chair)^1,2^, Ines Barroso (Deputy Chair)^3^, Jenefer M Blackwell^4^, ^5^, Elvira Bramon^6^, Matthew A Brown^7^, Juan P Casas^8^, Aiden Corvin^9^, Panos Deloukas^3^, Audrey Duncanson^10^, Janusz Jankowski^11^, Hugh S Markus^12^, Christopher G Mathew^13^, Colin NA Palmer^14^, Robert Plomin^15^, Anna Rautanen^1^, Stephen J Sawcer^16^, Richard C Trembath^13^, Ananth C Viswanathan^17^, Nicholas W Wood^18^

### Data and Analysis Group

Chris C A Spencer^1^, Gavin Band^1^, Céline Bellenguez^1^, Colin Freeman^1^, Garrett Hellenthal^1^, Eleni Giannoulatou^1^, Matti Pirinen^1^, Richard Pearson^1^, Amy Strange^1^, Zhan Su^1^, Damjan Vukcevic^1^, Peter Donnelly^1,2^

### DNA, Genotyping, Data QC and Informatics Group

Cordelia Langford^3^, Sarah E Hunt^3^, Sarah Edkins^3^, Rhian Gwilliam^3^, Hannah Blackburn^3^, Suzannah J Bumpstead^3^, Serge Dronov^3^, Matthew Gillman^3^, Emma Gray^3^, Naomi Hammond^3^, Alagurevathi Jayakumar^3^, Owen T McCann^3^, Jennifer Liddle^3^, Simon C Potter^3^, Radhi Ravindrarajah^3^, Michelle Ricketts^3^, Matthew Waller^3^, Paul Weston^3^, Sara Widaa^3^, Pamela Whittaker^3^, Ines Barroso^3^, Panos Deloukas^3^.

### Publications Committee

Christopher G Mathew (Chair)^13^, Jenefer M Blackwell^4,5^, Matthew A Brown^7^, Aiden Corvin^9^, Chris C A Spencer^1^

### Footnotes

1 Wellcome Trust Centre for Human Genetics, University of Oxford, Roosevelt Drive, Oxford OX3 7BN, UK;

2 Dept Statistics, University of Oxford, Oxford OX1 3TG, UK;

3 Wellcome Trust Sanger Institute, Wellcome Trust Genome Campus, Hinxton, Cambridge CB10 1SA, UK;

4 Telethon Institute for Child Health Research, Centre for Child Health Research, University of Western Australia, 100 Roberts Road, Subiaco, Western Australia 6008;

5 Cambridge Institute for Medical Research, University of Cambridge School of Clinical Medicine, Cambridge CB2 0XY, UK;

6 Department of Psychosis Studies, NIHR Biomedical Research Centre for Mental Health at the Institute of Psychiatry, King’s College London and The South London and Maudsley NHS Foundation Trust, Denmark Hill, London SE5 8AF, UK;

7 University of Queensland Diamantina Institute, Brisbane, Queensland, Australia;

8 Dept Epidemiology and Population Health, London School of Hygiene and Tropical Medicine, London WC1E 7HT and Dept Epidemiology and Public Health, University College London WC1E 6BT, UK;

9 Neuropsychiatric Genetics Research Group, Institute of Molecular Medicine, Trinity College Dublin, Dublin 2, Eire;

10 Molecular and Physiological Sciences, The Wellcome Trust, London NW1 2BE;

11 Department of Oncology, Old Road Campus, University of Oxford, Oxford OX3 7DQ, UK, Digestive Diseases Centre, Leicester Royal Infirmary, Leicester LE7 7HH, UK and Centre for Digestive Diseases, Queen Mary University of London, London E1 2AD, UK;

12 Clinical Neurosciences, St George’s University of London, London SW17 0RE;

13 King’s College London Dept Medical and Molecular Genetics, King’s Health Partners, Guy’s Hospital, London SE1 9RT, UK;

14 Biomedical Research Centre, Ninewells Hospital and Medical School, Dundee DD1 9SY, UK;

15 King’s College London Social, Genetic and Developmental Psychiatry Centre, Institute of Psychiatry, Denmark Hill, London SE5 8AF, UK;

16 University of Cambridge Dept Clinical Neurosciences, Addenbrooke’s Hospital, Cambridge CB2 0QQ, UK;

17 NIHR Biomedical Research Centre for Ophthalmology, Moorfields Eye Hospital NHS Foundation Trust and UCL Institute of Ophthalmology, London EC1V 2PD, UK;

18 Dept Molecular Neuroscience, Institute of Neurology, Queen Square, London WC1N 3BG, UK.

